# Land surface temperature evolution in rapidly urbanizing areas of Southeast Asia: studies from Vietnam and Cambodia

**DOI:** 10.1101/2025.07.09.663828

**Authors:** Bijeesh Kozhikkodan Veettil, Juliana Costi, Vanna Teck, Vikram Puri

**Affiliations:** Laboratory of Ecology and Environmental Management, Science and Technology Advanced Institute, Van Lang University, Ho Chi Minh City, Vietnam; Faculty of Applied Technology, School of Technology, Van Lang University, Ho Chi Minh City, Vietnam; Instituto de Oceanografia, Universiade Federal do Rio Grande, Rio Grande, Brazil; Faculty of Science and Technology, Svay Rieng University, Svay Rieng City, Cambodia; School of Computer Science, Duy Tan University, Da Nang, Vietnam; Institute of Research and Development, Duy Tan University, Da Nang, Vietnam

**Keywords:** Land surface temperatures, Urban microclimate, Urban heat island, Urban remote sensing, Urban sprawl

## Abstract

Land surface temperature (LST) is one of the crucial variables in urban microclimate studies. Satellite-based thermal data and vegetation indices, like the normalized difference vegetation index (NDVI), offer valuable insights into changes in LST and the development of urban heat islands (UHI). We analyzed the variations in LST and vegetation coverage in two rapidly urbanizing provinces, each in Southern Vietnam and Cambodia, respectively, during the 10 years between 2013 and 2024. In addition, complementary ERA5 Interim air temperature data were also used. Spatiotemporal changes in NDVI showed rapid urbanization in the eastern region of Battambang city and throughout the southern areas of Binh Duong Province. Time-series analysis indicated a consistent increase in LST in both study sites. There has been a notable increase in minimum LST since 2017 in the entire city of Battambang, whereas the central area of Battambang has become consistently warmer after 2020. The LST in southern Binh Duong gradually increased between 2014 and 2025 due to rapid urbanization and vegetation loss. The outcome of this study holds considerable importance, as the phenomenon of UHI formation has been documented in rapidly expanding cities globally, especially in Southeast Asia.

## 1. Introduction

The physics of land surface processes is highly influenced by land surface temperatures (LST) (Li et al. 2013). LST is a crucial parameter in the study of urbanization and climate change (Hereher 2016). Several studies (Tayyebi et al. 2018; Veettil et al. 2023) have analyzed the connection between land use changes (rapid urbanization in particular) and LST variations. Urban structures with higher heat storage capacity and decreased evaporation in cities lead to higher minimum land surface temperatures (LST) (Argüeso et al. 2014). Urbanization impacts on LST attracted the scientific community as it is believed to influence the local climate in cities compared to its neighborhood (Zeleňáková et al. 2015). These variations in microclimate between urban areas and their surroundings have been studied based on air temperature (Ojeh et al. 2016) or LST (Peng et al. 2018) measurements. LST data provides a reliable measure for assessing urban microclimate patterns (Fonseka et al. 2019). Satellite-based thermal infrared imagery has been used for LST retrieval by applying radiative transfer equations since the 1970s (Li et al. 2013).

Global urbanization has substantially impacted environmental and climate systems, intensifying climate change effects (Bazrkar et al. et al. 2015). For this reason, urbanization has attracted environmental scientists in the 21^st^ century (Veettil et al. 2023). Urbanization is a key anthropogenic activity that negatively impacts ecosystems, biodiversity, and urban microclimate (Guo et al. 2012; Veettil and Grondona 2018). Rapid urban growth lacking adequate planning is anticipated to aggravate these harmful consequences (Ranagalage et al. 2014). Therefore, a recent study by Maheshwari et al. (2020) suggested considering urbanization as one of the parameters in developing climate change adaptation strategies and policies to deal with future climate change scenarios.

UHI effect is characterized by increased air and ground temperatures in cities as compared to adjacent rural areas (Voogt and Oke, 2003), has been studied by scientists worldwide in the past few decades. Large cities and areas undergoing rapid urbanization may experience higher temperatures compared to rural regions because of the formation of UHI (Van and Bao 2020). The magnitude and geographic concentration of UHI-associated risks can be evaluated using UHI intensity and spatial distribution data (Cetin et al. 2024). Urban land cover types significantly affect UHI formation. Land cover modifications can be evaluated through the association between LST and vegetation indices like the Normalized Difference Vegetation Index (NDVI) (Singh and Grover 2015). Therefore, the formation and development of UHI can be analyzed by measuring LST measurements using satellite or unmanned aerial vehicle (UAV) imagery with thermal infrared information, which can be used as input for eco-friendly urban planning and development initiatives (Veettil et al. 2023).

Time-series LST values from remote sensing are considered as a proxy measurement in understanding the formation of UHIs (e.g., Rao 1972; Son et al. 2017; Veettil and Grondona 2018; Zhou et al. 2019; Veettil et al. 2023; Veettil and Van 2023). Satellite data with thermal infrared wavelengths, such as the Landsat series (except Multispectral Scanner-MSS) and ASTER, have proven their capability in measuring LST from space (e.g., Veettil and Grondona 2018; Veettil et al. 2023). Different computational approaches, such as automatic, semiautomatic, and machine learning algorithms, have been established for determining LST and UHI intensity. In addition to rapid measurements, satellite data offers cost-effective solutions for measuring LST compared to field measurements, as logistic expenses are high in the latter (Veettil et al. 2023; Veettil and Van 2023), particularly when time-series analysis is required. The availability of thermal infrared in the formation of satellite data has revolutionized UHI research (Tomlinson et al. 2011; Veettil et al. 2023).

The spatiotemporal variations in land surface temperatures in two rapidly urbanizing provinces, each from Vietnam and Cambodia, were analyzed during the period between 2013-2025 using satellite data having thermal infrared imagery. This analysis offers a comprehensive insight into the effect of alterations in land use and land cover (LULC) on the formation of UHIs and the potential consequences of other human-induced processes on the development of UHIs.

## 2. Study area

It is well known that industrialization and urbanization have reduced poverty in rural areas of Vietnam (Arouri et al. 2017). Such areas that underwent rapid urbanization were previously dependent on agriculture for income, and later (since 2010) these land areas were acquired by the government for urbanization for the socioeconomic transition in Vietnam (Thi et al. 2020). The present study considered two rapidly urbanizing provinces from Vietnam and Cambodia, Binh Duong and Battambang, respectively (**Figure 1**). The quality of environmental elements such as soil, water, and air has worsened in large urban and semi-urban areas over the past few decades because of rapid urbanization (Veettil and Van 2023; Veettil et al. 2023).

**Figure 1:**
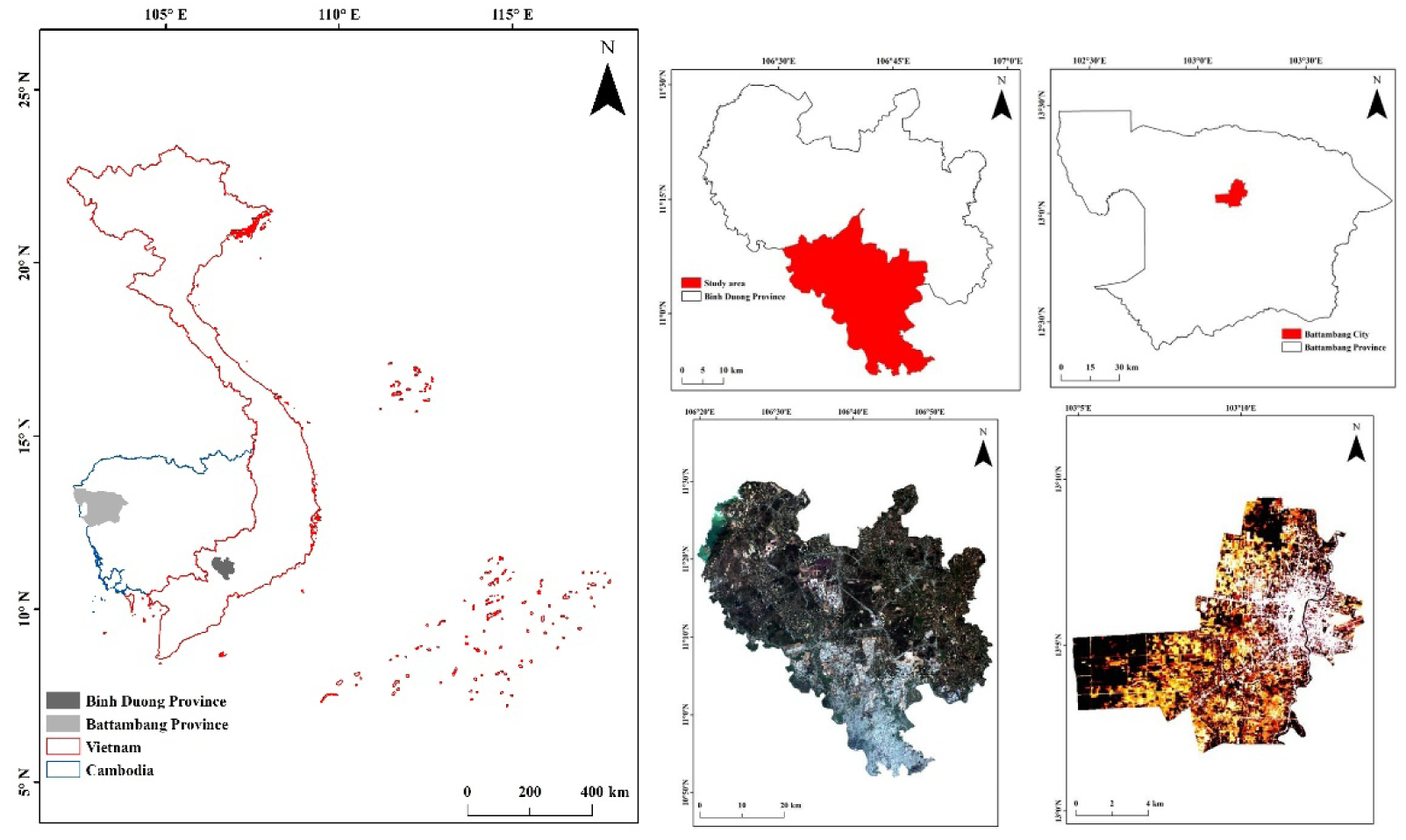
Study sites considered in Vietnam and Cambodia

Several studies conducted in Vietnam, including those in Hanoi and Ho Chi Minh City, observed the formation and development of UHI effect (Hoan et al. 2018; Son et al. 2017; Van and Bao 2020; Veettil and Van 2023), and small urbanized areas, such as Danang and Hue (Veettil et al. 2023). Son et al. (2017) reported that the LST values in Ho Chi Minh City changed from 31 °C in the 1990s to 32°C in the 2010s. Veettil and Van (2023) found a modest diminution in LST values in Ho Chi Minh City during the prolonged lockdown measures implemented because of COVID-19.

In the recent study, LST patterns in two rapidly urbanizing provinces, each from Vietnam and Cambodia, were analyzed during the period between 2014-2024. For this purpose, two urban areas were considered for estimating LST variations during the study period from satellite imagery. In addition to LST variations, this study also considered the vegetation dynamics in the study areas during the same period by using vegetation indices.

### 2.1. Binh Duong, Vietnam

Binh Duong Province, spanning an area of 2,694 square kilometers, is situated in southeastern Vietnam, renowned for its industrial significance. Thu Dau Mot City (10°58′N, 106°39′E) is the capital of Binh Duong Province in southeastern Vietnam. The advancement of industries has significantly added to the province’s economic growth. In parallel with the expansion of the urbanized areas and population growth, the land cover and land use in this region also changed significantly over the past three decades. It is worth noting that 77% of the land cover is used for agriculture (mostly rubber and fruit trees) and only 4% is the natural forest cover (Thuy et al. 2021).

The climate conditions in Binh Duong are influenced by tropical monsoons with an average annual temperature and rainfall of 26.5 °C and 1900 mm, respectively, which are suitable for rich vegetation diversity as the soil in this region is fertile (BDPC 2018). Additionally, the province contains 3 major rivers (Saigon River, Dong Nai River, and Song Be River) and multiple canals.

From 1995 to 2020, Binh Duong Province experienced a significant shift in land use, transitioning from agricultural and unused areas to various other types of land use, as observed by Bui and Mucsi (2022). This was due to the expansion of urbanized areas, mining sites, and artificially irrigated areas into land areas previously used for annual crops or unused croplands. The same study (Bui and Mucsi 2022) also showed that the expansion of urbanized areas in the province was 65 times between 1995 and 2020. Nevertheless, the root cause of rapid urbanization in the province is the high industrialization rate. Moreover, a recent study by Khuc et al. (2023) presented that the concentration of fine particles in the atmosphere in Binh Duong Province has increased in proportion to the expansion of urbanized regions and the concentration of people. Urban flooding is one of the negative effects of infrastructure development without proper planning, and few studies have suggested the implementation of a green infrastructure strategy for managing urban flooding in Binh Duong (An and Huong 2023). Nevertheless, the increase in LST due to rapid urban growth in this rapidly growing metropolitan area is relevant to investigate. The present study considered five highly urbanized districts in southern Binh Duong, namely Thu Dau Mot City, Di An, Thuan An, Ben Cat, and Tan Uyen.

### 2.2. Battambang, Cambodia

Battambang City, situated at 13°06′N and 103°12′E, is the capital of Battambang Province. It ranks as Cambodia’s second-largest city and is situated in the fifth-largest province of the country. Positioned in northwestern Cambodia, the city is approximately 300 kilometers from the national capital, Phnom Penh. According to the Department of Planning (2017), Battambang City had a population of 161,072 residents in 2016. The city encompasses a total land area of 115.44 km², with built-up areas constituting 25% of this territory (Battambang Municipality 2015). Agriculture is the primary economic activity in Battambang Province. Carter et al. (2016) describe the province as "the rice bowl of Cambodia," underscoring its significance in the nation’s agricultural sector.

Battambang experiences a tropical monsoon climate characterized by two distinct seasons: the rainy season, spanning from May to October, and the dry season, extending from November to April (Sourn et al. 2021). The average yearly temperature is 27.7°C, and the area gets about 1,331 mm of rain each year (Nut et al. 2021).

Han and Lim (2019) state that Battambang City lacked an official spatial development plan until 2015. Subsequent to this period, the national government sanctioned the "Land Use Master Plan of Battambang City, 2030." This Master Plan delineates the spatial organization of the city and outlines the planned long-term purposes of different land-use areas within the governing region. The strategy encompasses comprehensive maps that encompass different sections, including land-utilization zones, transit networks, regions for safeguarding cultural heritage, and initiatives for budget-friendly housing (Han and Lim 2019).

It is seen that there has been a rapid growth in the population of Battambang City since 2008. While there was only a 0.8% annual growth in population between 2008-2015, it has changed to 5.2% between 2015-2016 (Department of Planning 2017; Han and Lim 2019). Research conducted by Sourn et al. (2022) revealed that farmland in Battambang province expanded from 44.50% in 1998 to 61.11% in 2008 and continued to grow to 68.40% in 2018. At the same time, the same study estimated that the forest coverage dropped from 29.82% to 6.18% between 1998 and 2018. In addition, Sourn et al. (2021) estimated an increase of 4600 ha in built-up areas in Battambang Province, particularly in cities, between 2003 and 2008. Consequently, there will be an increase in the demand for housing and essential urban infrastructure, including roads, drainage and sewage networks, and freshwater provision (Han and Lim 2019). Nevertheless, the increase in LST due to rapid urban growth in this rapidly growing metropolitan area is also relevant to investigate.

## 3. Data and Methods

LST data, obtained from thermal infrared wavelengths in satellite imagery, has been utilized as a substitute for studying UHIs since the 1970s (Rao 1972). Since the launch of Landsat TM imagery in the 1980s, Landsat TM imagery has been used to monitor land use and land cover changes, followed by ASTER imagery, both have thermal data, and several studies on UHI formation expansion based on LST variations have been conducted (Zhou et al. 2019; Veettil et al. 2023). LST is regarded as a crucial element for determining the Earth’s surface’s radiative burden (de Almeida et al. 2021). As the LST patterns vary from small-scale to large-scale urban areas, LST information can be helpful in the study of urban microclimate (Heinl et al. 2015). Fonseka et al. (2019) noted that the number of developed areas is positively associated with the increase in LST and negatively correlated with other regions, such as vegetation and water. Hully et al. (2019) explain that LST can be determined by calculating the emitted surface radiance. This involves correcting sensor radiance data for atmospheric effects and using the inverse Planck function, while considering changes in emissivity.

Despite the advancements in remote sensing technology since the early 1980s, the capabilities of thermal data derived from satellite sensors have been utilized in Vietnam only since the 2010s (Veettil et al. 2023). Earliest studies, such as Son et al. (2017) and Hoan et al. (2018) studied UHI developments in Ho Chi Minh City in the south and Hanoi in the north, respectively, using Landsat imagery. Van and Bao (2020) used ASTER data instead of Landsat imagery for LST calculation in Ho Chi Minh City.

The current study employed Landsat-8 OLI and TIRS data acquired during the period between 2013 and 2025 to assess the fluctuations in LST in Binh Duong Province in Vietnam and Battambang Province in Cambodia. Landsat-8 OLI imagery possesses a spatial resolution of 30m, whereas that of TIRS images is 100m, in addition to the panchromatic imagery with a 15m resolution. Landsat series data can be accessed for free from the United States Geological Survey (USGS) via https://earthexplorer.usgs.gov/. This study utilized only cloud-free, Level 2 Landsat images that were acquired during the study period. Table 1 presents the detailed specifications of the satellite data utilized for each study site.

**Table 1:**
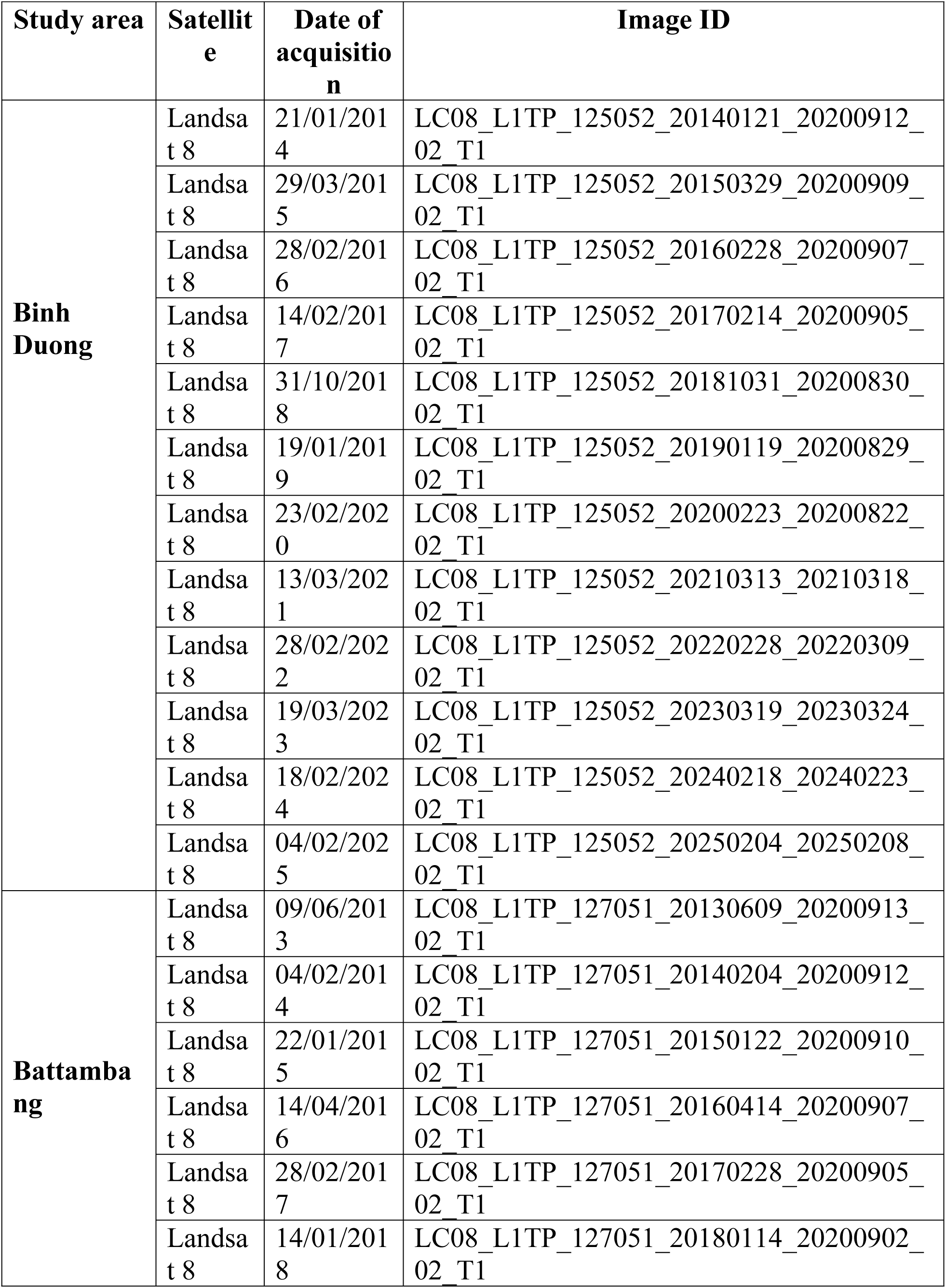
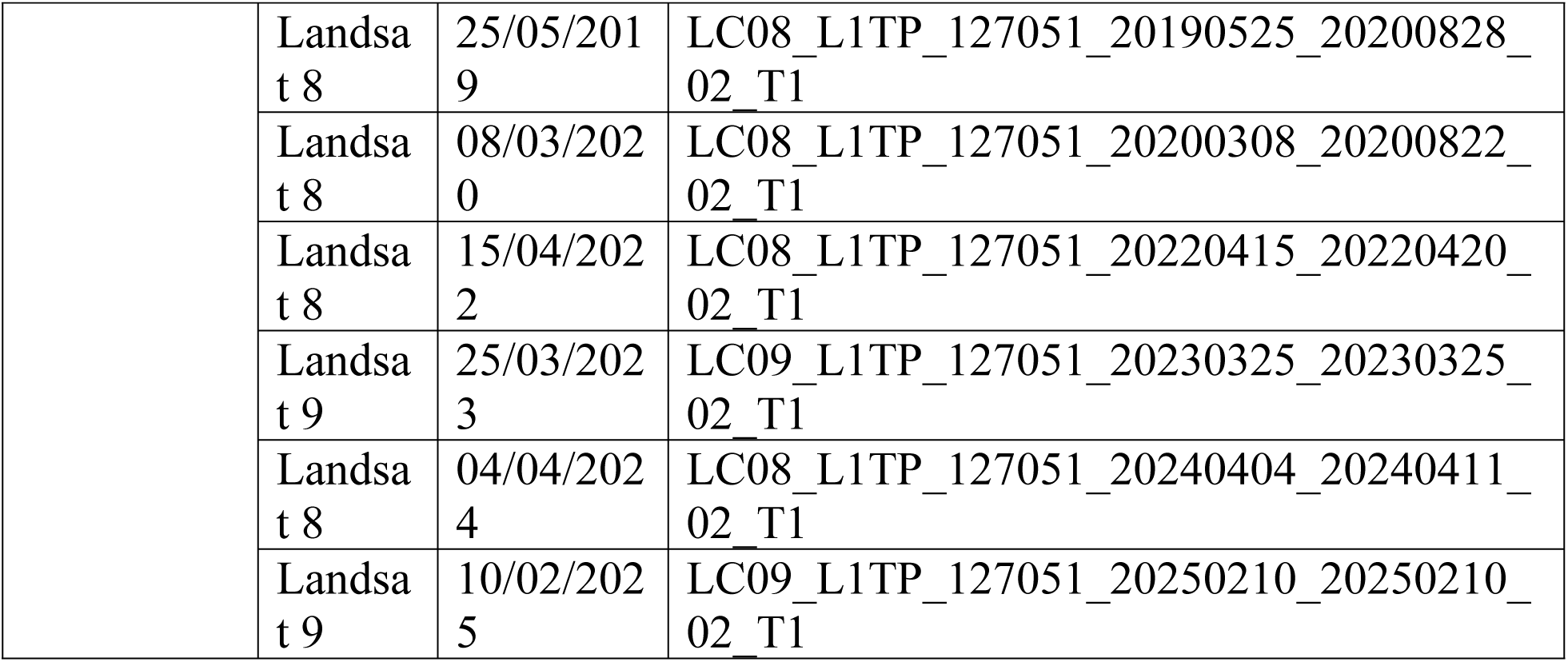
Landsat-8 and Landsat-9 data used in the study.

| Study area | Satellite | Date of acquisition | Image ID |
| --- | --- | --- | --- |
| **Binh Duong** | Landsat 8 | 21/01/2014 | LC08_L1TP_125052_20140121_20200912_02_T1 |
|  | Landsat 8 | 29/03/2015 | LC08_L1TP_125052_20150329_20200909_02_T1 |
|  | Landsat 8 | 28/02/2016 | LC08_L1TP_125052_20160228_20200907_02_T1 |
|  | Landsat 8 | 14/02/2017 | LC08_L1TP_125052_20170214_20200905_02_T1 |
|  | Landsat 8 | 31/10/2018 | LC08_L1TP_125052_20181031_20200830_02_T1 |
|  | Landsat 8 | 19/01/2019 | LC08_L1TP_125052_20190119_20200829_02_T1 |
|  | Landsat 8 | 23/02/2020 | LC08_L1TP_125052_20200223_20200822_02_T1 |
|  | Landsat 8 | 13/03/2021 | LC08_L1TP_125052_20210313_20210318_02_T1 |
|  | Landsat 8 | 28/02/2022 | LC08_L1TP_125052_20220228_20220309_02_T1 |
|  | Landsat 8 | 19/03/2023 | LC08_L1TP_125052_20230319_20230324_02_T1 |
|  | Landsat 8 | 18/02/2024 | LC08_L1TP_125052_20240218_20240223_02_T1 |
|  | Landsat 8 | 04/02/2025 | LC08_L1TP_125052_20250204_20250208_02_T1 |
| **Battambang** | Landsat 8 | 09/06/2013 | LC08_L1TP_127051_20130609_20200913_02_T1 |
|  | Landsat 8 | 04/02/2014 | LC08_L1TP_127051_20140204_20200912_02_T1 |
|  | Landsat 8 | 22/01/2015 | LC08_L1TP_127051_20150122_20200910_02_T1 |
|  | Landsat 8 | 14/04/2016 | LC08_L1TP_127051_20160414_20200907_02_T1 |
|  | Landsat 8 | 28/02/2017 | LC08_L1TP_127051_20170228_20200905_02_T1 |
|  | Landsat 8 | 14/01/2018 | LC08_L1TP_127051_20180114_20200902_02_T1 |
|  | Landsat 8 | 25/05/2019 | LC08_L1TP_127051_20190525_20200828_02_T1 |
|  | Landsat 8 | 08/03/2020 | LC08_L1TP_127051_20200308_20200822_02_T1 |
|  | Landsat 8 | 15/04/2022 | LC08_L1TP_127051_20220415_20220420_02_T1 |
|  | Landsat 9 | 25/03/2023 | LC09_L1TP_127051_20230325_20230325_02_T1 |
|  | Landsat 8 | 04/04/2024 | LC08_L1TP_127051_20240404_20240411_02_T1 |
|  | Landsat 9 | 10/02/2025 | LC09_L1TP_127051_20250210_20250210_02_T1 |

To complement the satellite-based LST analysis, ERA5 reanalysis data were used to assess temporal and spatial variations in near-surface air temperature (2 m) in Battambang (Cambodia) and Binh Duong (Vietnam) from 2013 to 2025. These datasets offer continuous hourly records with a spatial resolution of approximately 31 km × 31 km.

LST variations between 2013 and 2024 in the two study sites (Binh Duong-Vietnam and Battambang-Cambodia) were approximated using the Landsat-8 TIRS data and NDVI from Landsat-8 OLI data as described in Avdan and Jovanovska (2016). Several studies (Guha and Govil 2020) have discussed the relationship between LST and seasonal variations in NDVI. The 6-step algorithm used for estimating LST in °C (Avdan and Jovanovska 2016; Veettil et al. 2023; Veettil and Van 2023) is discussed below:

### a. Conversion of thermal infrared (TIR) digital number (DN) values into Top of the Atmosphere (TOA) spectral radiance

To begin estimating LST, Band 10 from the TIRS is transformed from Digital Number (DN) values to spectral radiance (R) at the top of the atmosphere using equation 1.

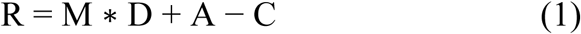

Where:

M = band-specific multiplicative rescaling coefficient;
D = Band 10 digital number values of the image;
A = band-specific additive rescaling coefficient;
C = correction factor for Band 10.

The values for M and A are located within the metadata of the Landsat-8 data package. Montanaro et al. (2014) advised the utilization of Band 10 TIRS data for the estimation of LST as opposed to Band 11, citing concerns regarding the calibration uncertainty of the latter.

### b. Conversion of TOA spectral reflectance into at-sensor brightness temperature (BT) in °C

The second step requires converting the TOA spectral radiance into brightness temperature (T) by applying equation 2.

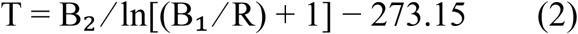

Where:

B₁ = band-specific thermal conversion coefficient 1 = 774.8853 W/(m².sr.μm);
B₂ = band-specific thermal conversion coefficient 2 = 1321.0789 K

B₁ and B₂ can be found in the metadata of the Landsat-8 data package. To convert these values to brightness temperature in °C, subtract 273.15.

### c. Normalized Difference Vegetation Index - NDVI

Many studies recognized that LST and land surface characteristics are closely correlated (Guha and Govil 2020). Due to evaporative cooling, a negative correlation between NDVI and LST is expected (Deng et al. 2018). This negative correlation can be corrected by using NDVI to estimate LST, which can be used to estimate emissivity. NDVI can be estimated by using equation 3. NDVI is vital for determining land surface emissivity, which is necessary for calculating land surface temperature (Guha and Govil 2020).

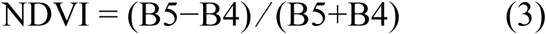

B4 = Landsat 8 Band 4 (near infrared)
B5 = Landsat 8 Band 5 (red)

Seasonal variations in the NDVI-LST relationship have been considered as an important variable in the study of mixed urban land surfaces (Guha and Govil 2020). Generally. The negative correlation between NDVI and LST is stronger in the wet season than the dry season (Guha et al. 2020). Some studies, like Ullah et al. (2023), found that NDVI increases with elevation. This information is important for high-altitude urban areas. This observation can be useful in regulating temperature in urban areas by applying a combination of vegetation and height variables, depending on the availability (Alexander 2021).

### d. Proportion of vegetation (Pv)

To use NDVI for estimating LST, it is crucial to know the amount of vegetation. Deardorff (1978) characterized Fv as the proportion of the ground area that is in contact with vegetation, including leaves, stems, and branches, relative to the entire area occupied by vegetation. The proportion of vegetation (equation 4) is a key biophysical variable related to earth surface processes, important for monitoring biodiversity and modeling climate and weather (Gutman and Ignatov 1998).

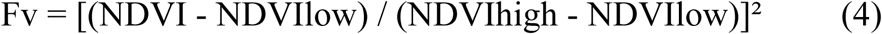

Where:

NDVIlow = the minimum value of NDVI;
NDVIhigh = the maximum value of NDVI.

### e. Estimation of emissivity (ε) from the Proportion of vegetation

After estimating the proportion of vegetation, the next step is to calculate the emissivity using equation 5. A higher proportion of vegetation leads to a higher overall emissivity for the surface.

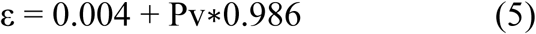

### f. Estimation of LST from brightness temperature and emissivity

Once you know the brightness temperature and emissivity, you can calculate the LST in degrees Celsius using equation 6.

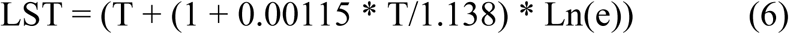

Where:

T = brightness temperature
e = emissivity

Air temperature at 2 meters above the surface was obtained from the ERA5 reanalysis, produced by the European Centre for Medium-Range Weather Forecasts (ECMWF). ERA5 provides global climate data at an hourly temporal resolution and a horizontal resolution of approximately 31 km × 31 km. The dataset was accessed for the period from 2013 to 2025, covering the spatial domain between 10.8°S and 13.05°S latitude and 102.8°E to 106.8°E longitude. For each year, hourly temperature values were spatially averaged within the boundaries of each province, as defined by the administrative shapefiles. Daily means were computed by averaging all hourly values for each complete day, followed by monthly and annual means derived from the daily time series. These temporal aggregations were used to characterize the seasonal and interannual variability in near-surface air temperature across the region. The ERA5 dataset is described in detail by Hersbach et al. (2020).

## 4. Results and Discussion

This study looked at changes in NDVI and LST in two provinces that are growing quickly due to urbanization, one in Cambodia and the other in Vietnam. The NDVI series helped to understand the spatiotemporal changes in land use, whereas the LST series has been employed to explain the influence of land use on urban microclimate.

### 4.1. Trends in NDVI

Typically, NDVI ranges from -1 to +1; a value near +0.6 indicates a healthy (forests or crops) vegetation, whereas a value near 0 indicates sparse vegetation or bare land, and negative NDVI values indicate water or non-vegetated surfaces. A temporal change in NDVI in agricultural areas indicates seasonal crop cycles, land use changes, or impacts of climate change (e.g., desertification). However, a gradual decrease in NDVI in a time-series map indicates urban sprawl, deforestation, or soil degradation.

The time-series map of NDVI in Binh Duong province also showed its fast-growing urban areas during the past decade (**Figure 2a**), particularly in its central and southern regions. The observed changes in NDVI agree with previous studies, such as Bui and Musci (2022), which estimated a 65-fold expansion of urban areas between 1995 and 2020 (25 years) in Binh Duong province. Another recent study by Huyen et al. (2022) also came with similar results on impervious surfaces. The same study (Huyen et al. 2022) highlighted the importance of green areas, including green corridors, natural forests, and planted forests. Compared to Battambang City, where urbanization is concentrated in the eastern areas, the entire southern Binh Duong province underwent urbanization during the study period.

**Figure 2:**
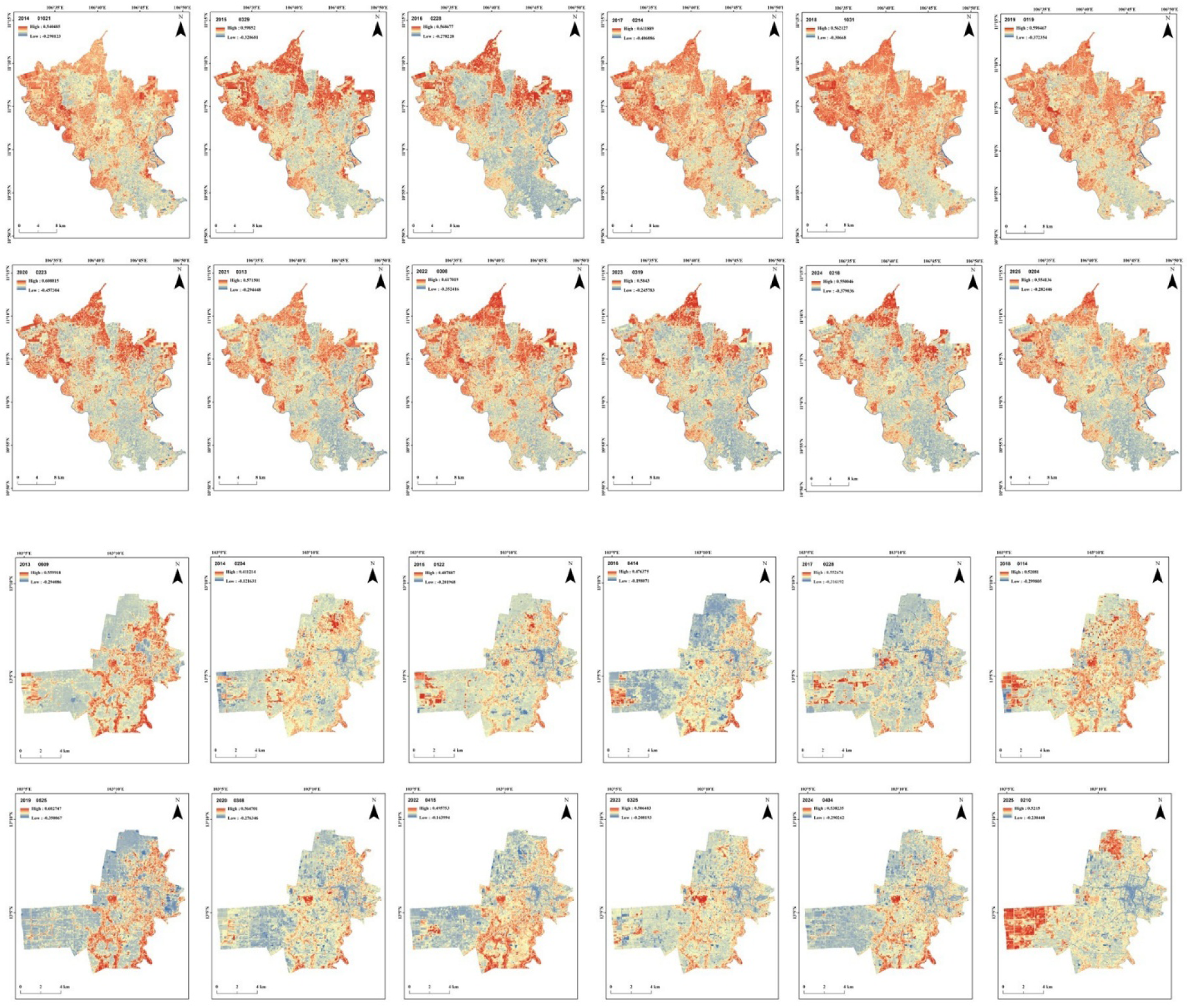
Spatiotemporal variations in NDVI in Binh Duong (top) and Battambang (bottom).

The changes in NDVI patterns over time and space in Battambang city demonstrate the influence of urban development on land use patterns transformations. The swift growth of metropolitan regions in the eastern regions of the study site is visible from the resulting time-series map of NDVI (**Figure 2b**). The differences in NDVI seen on the western side are due to seasonal crops and shifts in land use patterns. The rapid reduction in NDVI along the eastern side of the study area agrees with a previous study (Sourn et al. 2021) that estimated an exceptional increase in built-up areas from 48 ha to 4698 ha between 1998 and 2018. In contrast, the forest area has reduced from 358.96 hectares to 74.42 hectares over the same time. Some unusual readings in NDVI in 2023 and 2025 might suggest land being turned into farmland or minor cloud cover affecting the satellite data.

The NDVI is highly responsive to plant life cycles and is widely regarded as an excellent measure of vegetation coverage and plant growth. It is also used to study changes in vegetation over various areas and periods. Both the study sites have shown fast-growing urban areas, which make them an ideal location to study the development of urban microclimate. A disadvantage of using NDVI for time-series analysis is that it is typically not stationary, experiencing seasonal variations and both long-term and short-term fluctuations (Martínez and Gilabert 2009). The present study used images taken during the same season of the year to avoid the effects of seasonality in vegetation from the satellite data. NDVI is important ecologically because it shows changes in biomass. This makes it a valuable tool for understanding the complex relationship between climate change, vegetation distribution patterns, and groundwater availability for woody plants over large areas and time periods (Aguilar et al. 2012).

### 4.2. Trends in land surface temperatures

The spatiotemporal changes in LST indicated both a decline in vegetation coverage and a rise in impervious surfaces in both study areas. A color scale from light to dark orange has been used to indicate LST (darker areas indicate a higher LST compared to lighter ones). Urban and densely populated areas consistently exhibit higher LST values throughout the study period, likely due to urban heat island formation.

The LST series between 2014 and 2025 in Binh Duong (**Figure 3a**) indicated a gradual increase in land surface temperatures during the study period. Spatial patterns indicate a consistently higher LST in urban areas across the years considered, indicating the formation of UHIs. Even though there is a fluctuation in LST, the general trend is an increase in maximum LST, likely due to urbanization, vegetation loss, and climate change.

**Figure 3:**
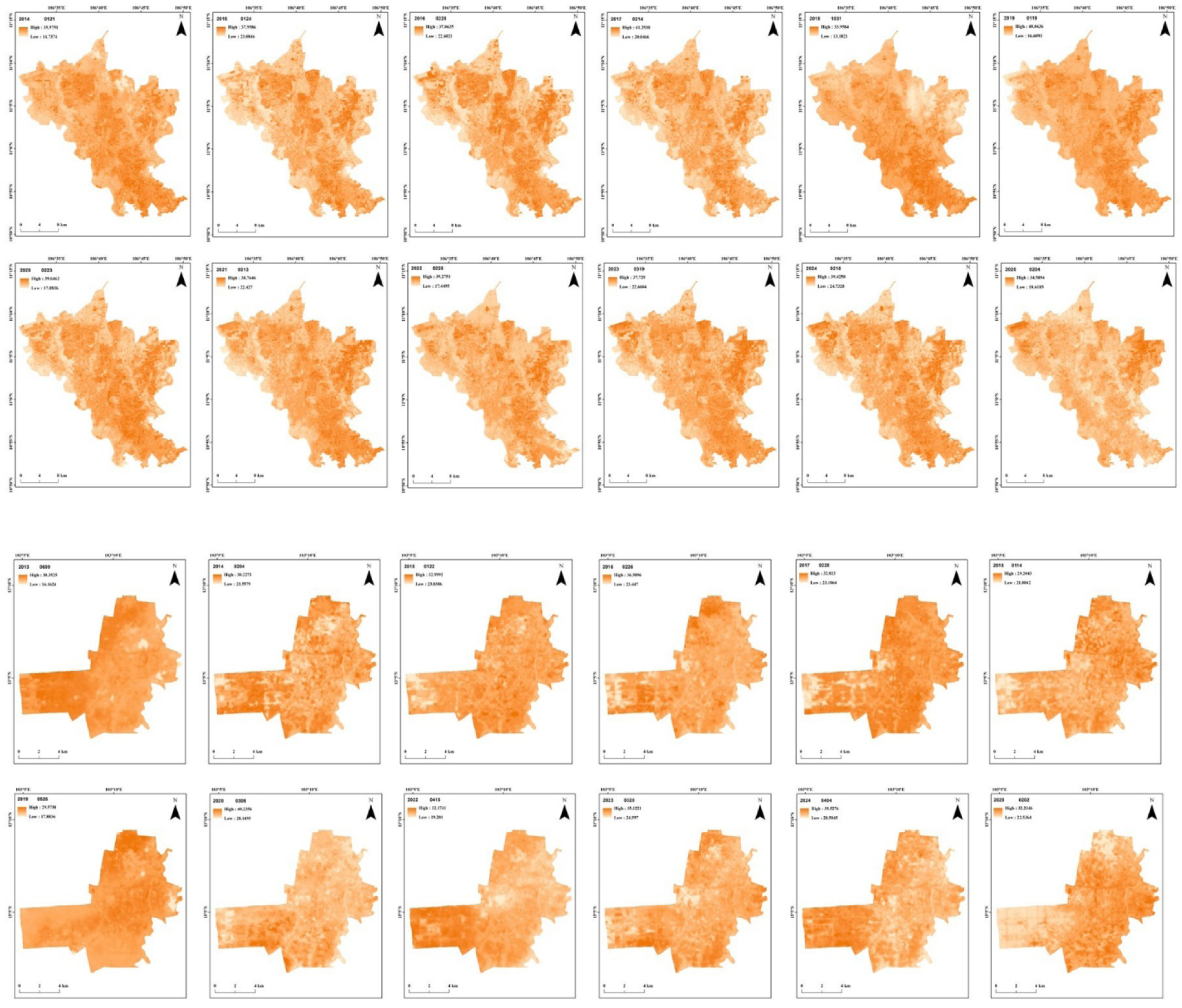
Spatiotemporal variations in land surface temperatures in Binh Duong (top) and Battambang (bottom).

The LST series in Battambang (**Figure 3b**) indicated a rise in LST, particularly after 2020, and the central region of the study area is consistently warmer due to the development of urban areas. There has been a notable increase in minimum LST since 2017, probably indicating warmer nights with less cooling. The LST range became narrow after 2020, indicating an overall warming with less variability.

### 4.3. Trends in near-surface air temperature (ERA5)

Time series of air temperature (**Figure 4**) reveal distinct seasonal cycles and interannual variability in both study sites. In general, the hourly and daily data show clear seasonal oscillations, while the monthly and annual aggregates provide insight into long-term patterns associated with urban development and land cover change.

**Figure 4:**
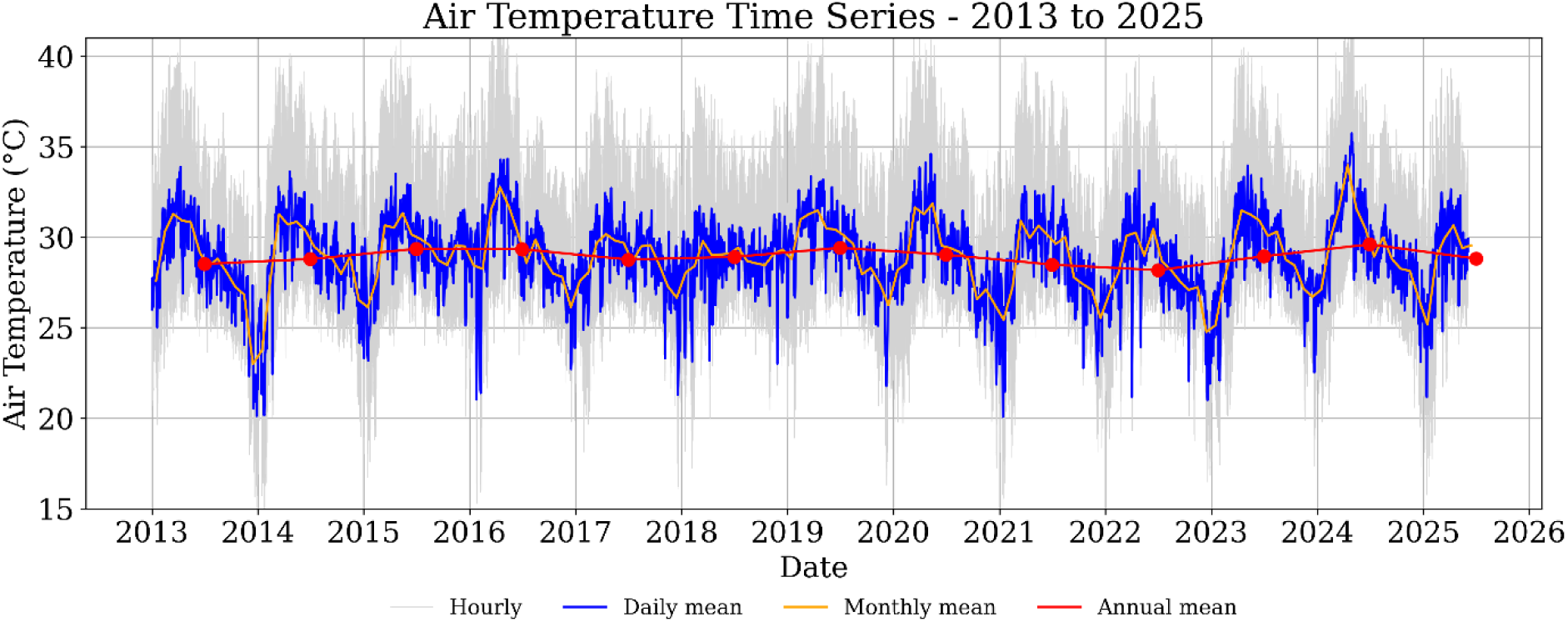
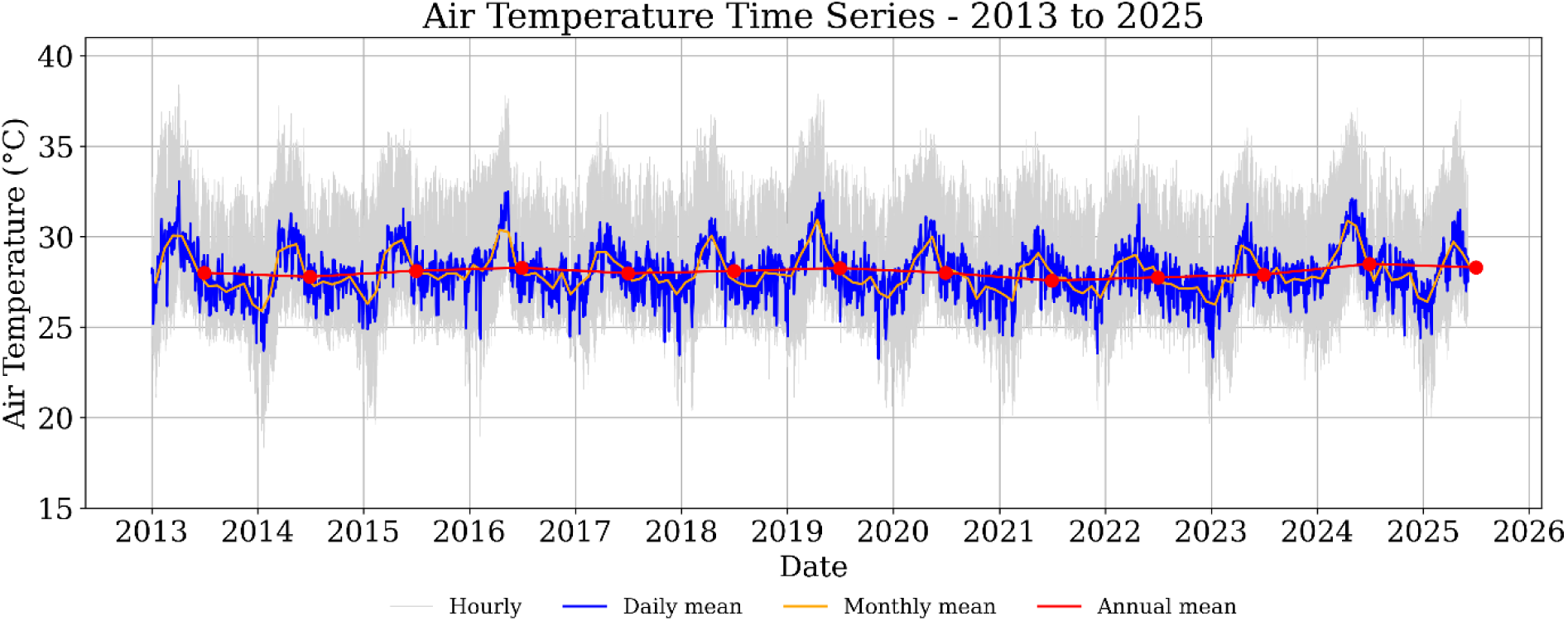
Time series of near-surface air temperature (2 m, ERA5) from 2013 to 2025 for Battambang (top) and Binh Duong (bottom). Hourly data are shown in gray, daily means in blue, monthly means in orange, and annual means in red with linear trend.

In Binh Duong, the annual mean air temperature remained relatively stable around 29 °C, with notable seasonal amplitude and limited variability in the long-term trend. The spatial maps of annual mean and maximum temperatures (**Figure 5** and **Figure 6**) show consistently higher values in the southern region of the province, especially in urban districts such as Thu Dau Mot, Di An, and Thuan An. These spatial patterns align with areas of intense urbanization and industrialization described in Section 2.1 and are indicative of the establishment of urban heat island (UHI) conditions. The maximum air temperature regularly exceeds 40 °C in some years, further supporting this hypothesis.

**Figure 5:**
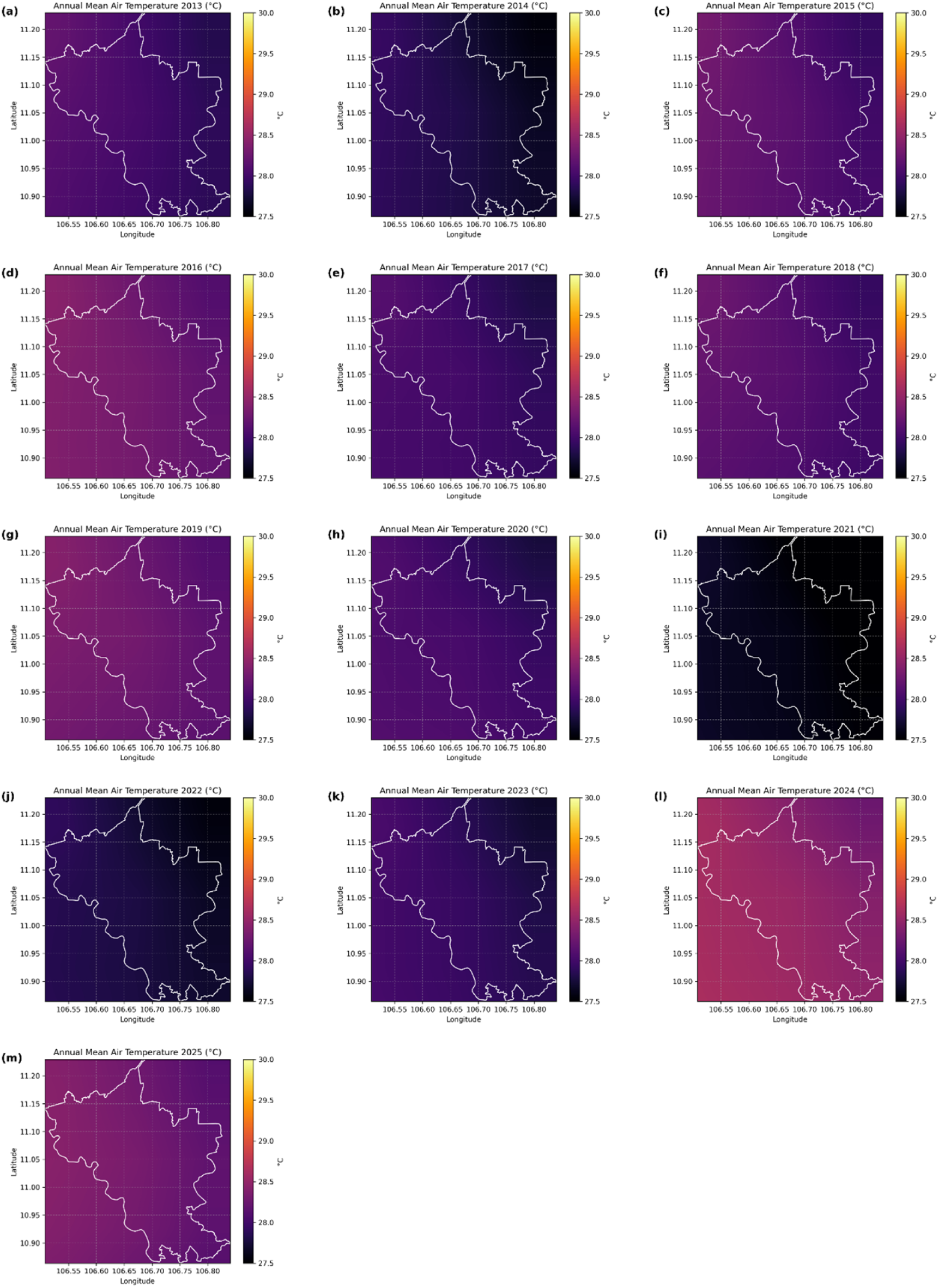
Annual mean air temperature (ERA5, °C) in Binh Duong, Vietnam, from 2013 to 2025. The white polygon shows the administrative boundary of the province.

**Figure 6:**
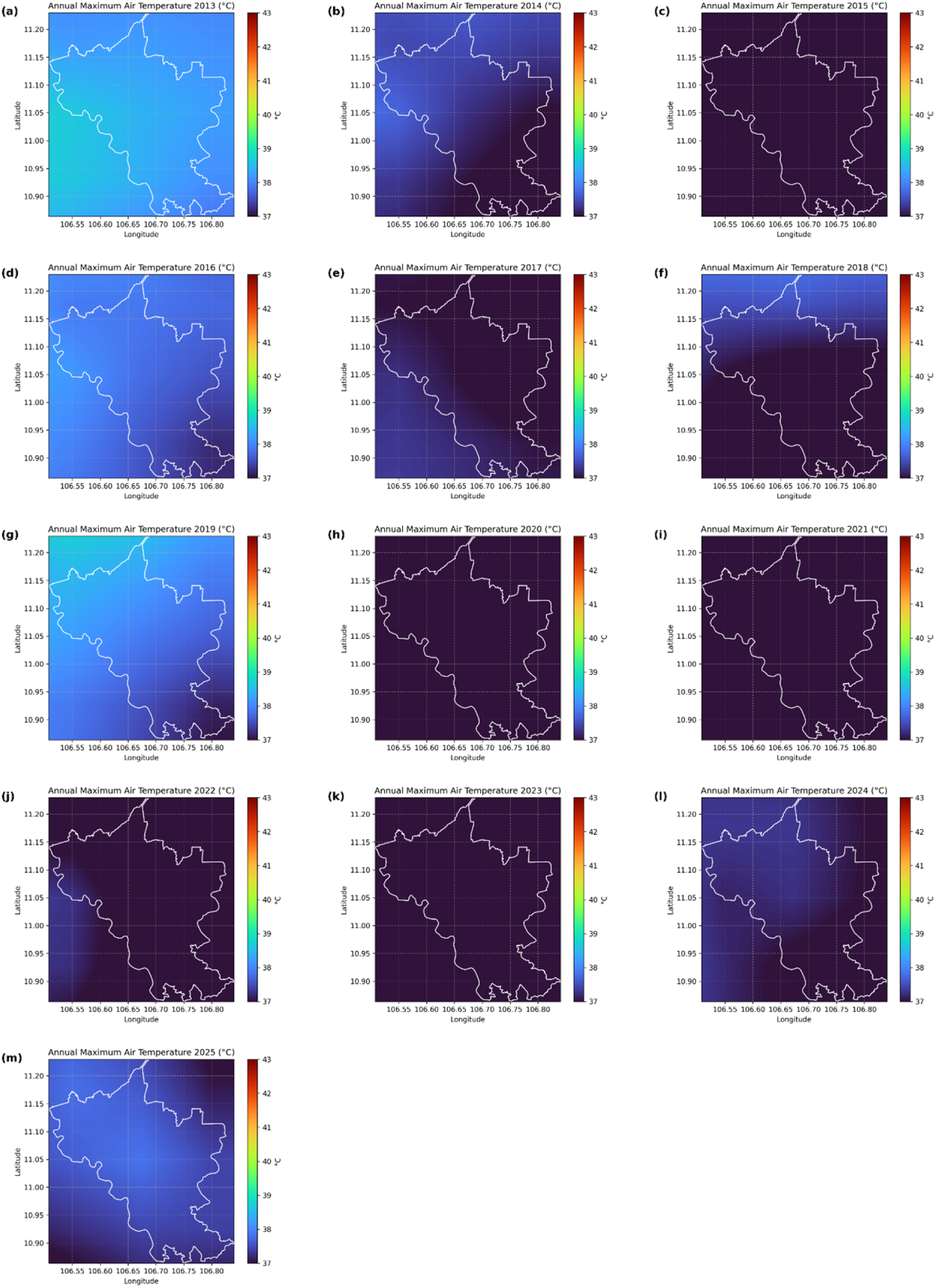
Annual maximum air temperature (ERA5, °C) in Binh Duong, Vietnam, from 2013 to 2025. The white polygon shows the administrative boundary of the province.

In contrast, Battambang Province exhibits a more pronounced temporal pattern, with increasing fluctuations in daily and monthly means observed particularly after 2020 (**Figure 4**). Despite interannual variability, the annual mean air temperature hovered around 28.5 °C during most of the period. Spatially, the maps of annual mean and maximum temperature (**Figure 7** and **Figure 8**) highlight persistent warming in the central and eastern parts of the province, regions undergoing accelerated urban development, as documented in Section 2.2. This observation is consistent with the reported increase in minimum LST in Battambang after 2017 and supports the inference of UHI intensification due to urban sprawl and vegetation loss.

**Figure 7:**
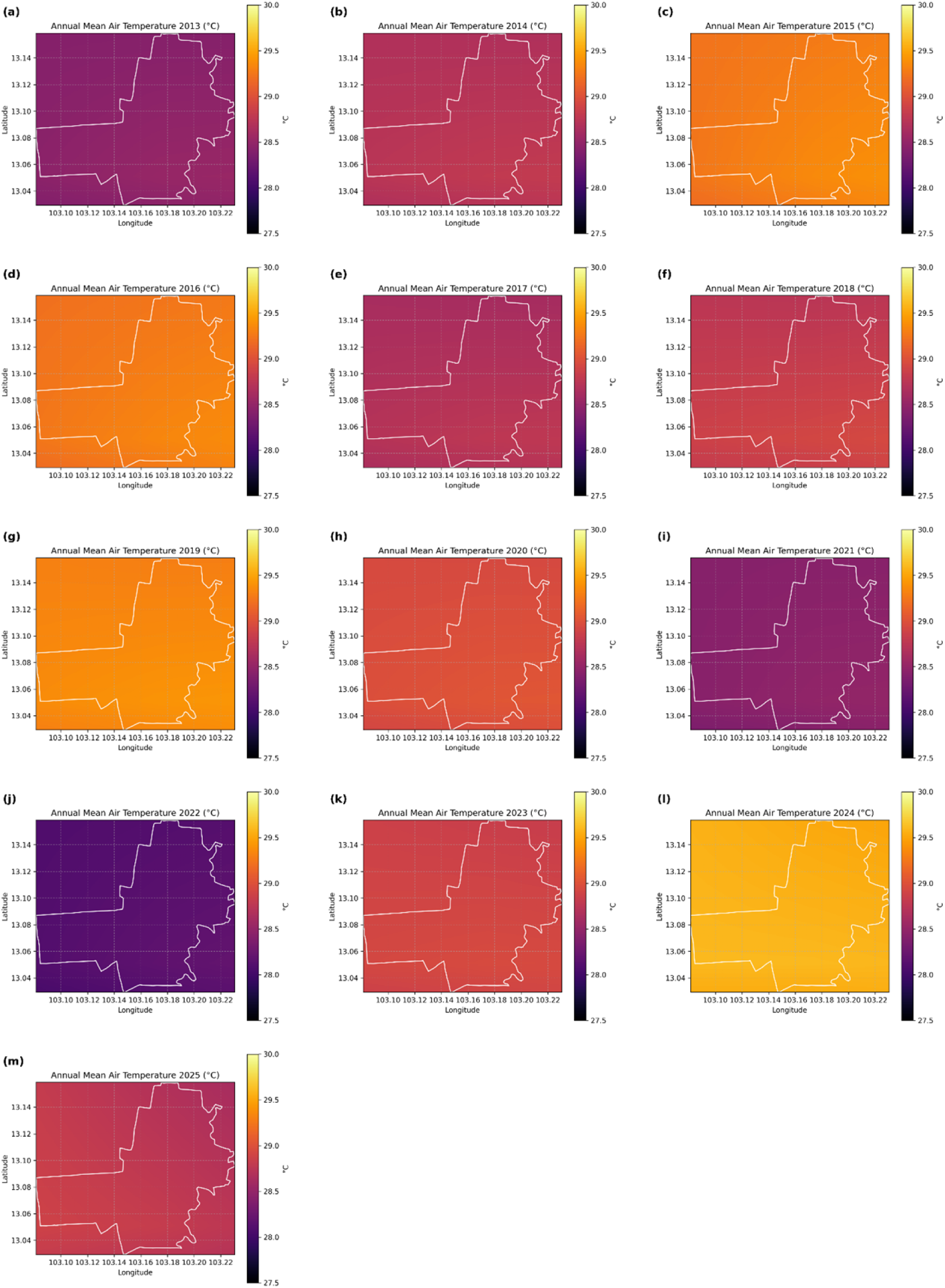
Annual mean air temperature (ERA5, °C) in Battambang, Cambodia, from 2013 to 2025. The white polygon shows the administrative boundary of the province.

**Figure 8:**
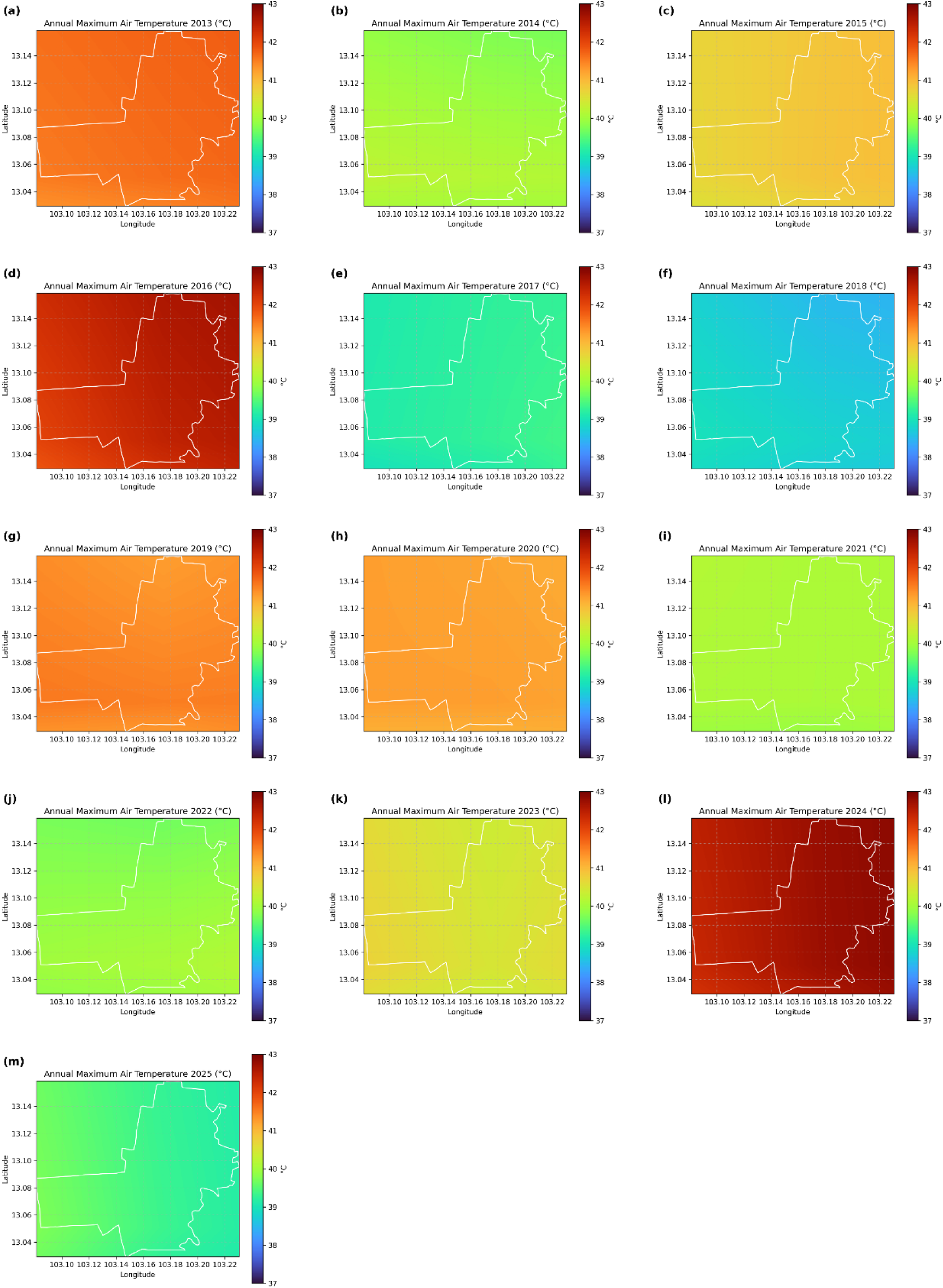
Annual maximum air temperature (ERA5, °C) in Battambang, Cambodia, from 2013 to 2025. The white polygon shows the administrative boundary of the province.

Overall, the analysis of ERA5 near-surface air temperature data corroborates the findings based on LST and NDVI, confirming that both study areas experienced localized warming trends attributable to urban expansion. The integration of reanalysis-based air temperature data provides an additional line of evidence for understanding the urban microclimate dynamics and enhances the robustness of UHI detection at the regional scale.

## 5. Conclusions

Fast urban growth leads to negative environmental effects, such as increased LST and the creation of UHI. These areas become warmer than their surroundings because they absorb sunlight and release it as heat. This study investigated the LST changes in two provinces undergoing swift urbanization, Vietnam (Binh Duong) and Cambodia (Battambang), between 2013 and 2025. LST variations are crucial for understanding how human activities affect urban areas, contribute to the creation of UHIs, and influence the urban microclimate.

The time-series analysis of NDVI indicated urban areas’ expansion and vegetation reduction in both Battambang and Binh Duong. While the rapid expansion of metropolitan areas in Battambang focused on its eastern region, the impervious surfaces have rapidly expanded in the entire southern Binh Duong province.

The spatiotemporal changes in LST indicated a decline in vegetation coverage and an increase in the area covered by impervious surfaces in Binh Duong and Battambang. The findings reveal that LSTs have risen over the years due to the creation of UHIs. These changes highlight the negative effects of quick urbanization and the growth of impervious surfaces, despite also contributing to financial stability and economic growth.

## Acknowledgments

We would like to acknowledge the United States Geological Survey (USGS) for providing Landsat imagery.

## Authors’ contributions

All authors contributed equally to the manuscript.

## Funding

No funding to declare.

## Data availability

Data will be available on request.

## Code availability

Not applicable.

## Declarations

### Competing interests

The authors declare no competing interests.

### Conflicts of interest/Competing interests

No conflicting/ competing interests to declare.

### Ethics approval/declarations

Not Applicable.

### Consent to participate

Not Applicable.

### Consent for publication

Not Applicable.

## Notes

### Competing Interest Statement

The authors have declared no competing interest.

